# Unmatched level of molecular convergence among deeply divergent complex multicellular fungi

**DOI:** 10.1101/549758

**Authors:** Zsolt Merényi, Arun N. Prasanna, Wang Zheng, Károly Kovács, Botond Hegedüs, Balázs Bálint, Balázs Papp, Jeffrey P. Townsend, László G. Nagy

## Abstract

Convergent evolution is pervasive in nature, but it is poorly understood how various constraints and natural selection limit the diversity of evolvable phenotypes. Here, we report that, despite >650 million years of divergence, the same genes have repeatedly been co-opted for the development of complex multicellularity in the two largest clades of fungi—the Ascomycota and Basidiomycota. Co-opted genes have undergone duplications in both clades, resulting in >81% convergence across shared multicellularity-related families. This convergence is coupled with a rich repertoire of multicellularity-related genes in ancestors that predate complex multicellular fungi, suggesting that the coding capacity of early fungal genomes was well suited for the repeated evolution of complex multicellularity. Our work suggests that evolution may be predictable not only when organisms are closely related or are under similar selection pressures, but also if the genome biases the potential evolutionary trajectories organisms can take, even across large phylogenetic distances.

## Introduction

Darwin suggested that organisms can evolve an unlimited variety of forms (*1*). Contrary to his concept of ‘unlimited forms’, it is now clear that evolution follows similar paths more often than classic models of genetic change would predict (*2*–*5*). The independent emergence of similar phenotypes is called convergent evolution, which happens in response to similar selection pressures, bias in the emergence of phenotypic variation (*6, 7*), or both (*3, 5, 8*). Convergence is widespread in nature (e.g. (*9*–*12*)) suggesting that evolution may be predictable (*13*) and deterministic (*5, 11, 14*) under some circumstances, although what drives divergent lineages to evolve similar phenotypes is poorly understood.

A fascinating example of convergent phenotypes is multicellularity: it has evolved at least 25–30 times across the pro-and eukaryotes (*15*–*23*), reaching a diversity of complexities that range from simple cell aggregates to the most complex macroscopic organisms (*24*). Instances of the evolution of multicellularity are considered major transitions in evolution—a conceptual label that is difficult to reconcile with repeated origins (*15, 17*). This difficulty follows from the assumption that major transitions are limited by big genetic hurdles and thus should occur rarely during evolution (*17, 25*).

The highest level of multicellular organization is referred to as complex multicellularity (CM), which, unlike unicells and simple multi-celled aggregates (e.g. filaments, colonies, biofilms, etc.), is characterized by a three-dimensional organization, sophisticated mechanisms for cell-cell adhesion and communication, and extensive cellular differentiation (*17, 23, 24, 26*). CM occurs in metazoans, embryophyte plants, and fungi as well as red and brown algae. In fungi, CM refers mostly to sexual fruiting bodies which are found in 8-11 disparate fungal clades and show clear signs of convergent origins (*22*). Although they originated independently, CM fungal clades are phylogenetically close, providing a tractable system for studying the genetics of major evolutionary transitions in complexity. Fruiting bodies in fungal lineages can be developmentally and morphologically highly distinct—yet they evolved for the same general purpose: to enclose sexual reproductive structures in a protective environment and facilitate spore dispersal (*23, 27*–*30*). Here we seek to explain the convergent evolution of fungal fruiting bodies by analyzing the fate of multicellularity-related gene families across the two largest clades of CM fungi: the Agaricomycotina (mushroom-forming fungi, Basidiomycota) and the Pezizomycotina (Ascomycota).

## Results

To study the evolution of complex multicellularity in fungi we first reconstructed ancestral cellularity levels across a phylogenetic tree of 19 representative species (table S1). This analysis strongly suggests that the two most diverse CM clades, Agarico-and Pezizomycotina, acquired fruiting body formation independently (Fig. 1/a). Likelihood proportions imply that the most recent common ancestor (MRCA) of the Dikarya did not form fruiting bodies (marginal probability of non-CM: 0.87, CM: 0.13), followed by two independent acquisitions of CM in the MRCA of the Agaricomycetes and MRCA of Pezizomycotina. Our data suggest that the MRCA of the Ascomycota did not produce fruiting bodies (state non-CM: 0.616, state CM: 0.384), which implies a third independent origin of fruiting body production in *Neolecta* (Taphrinomycotina) (*22, 31, 32*) (Fig. 1/a).

**Figure 1.**
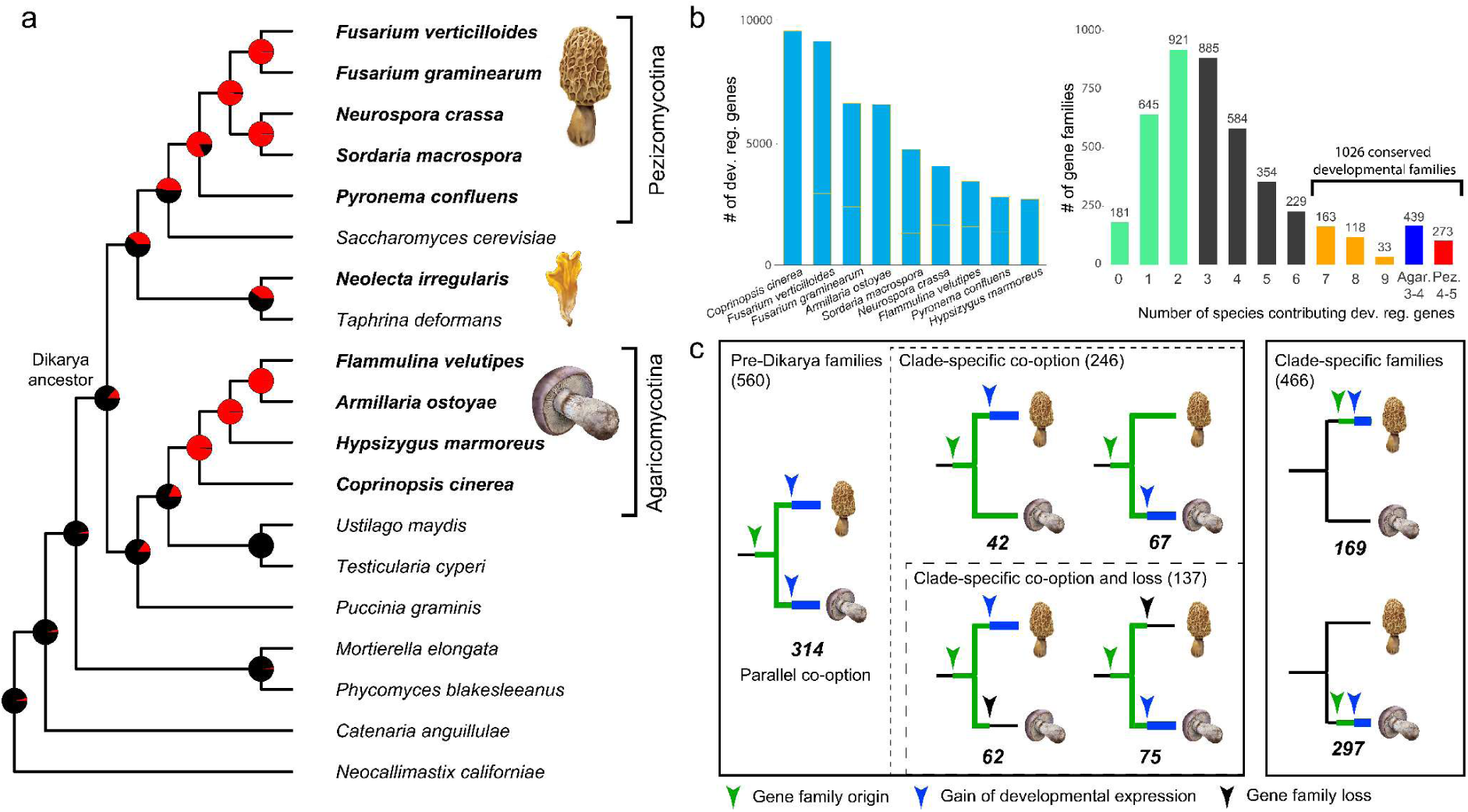
The evolution of complex multicellularity in fungi and conserved developmentally regulated gene families. **(a)** phylogenetic relationships among 19 species analyzed in this study inferred from 86 conserved, single-copy orthologs. Two independent clades of complex multicellular species are marked, and typical fruiting body morphologies are shown. Pie charts at nodes indicate the proportional likelihoods of CM (red) and non-CM (black) ancestral states reconstructed using Maximum Likelihood. Character state coding of extant species are shown as bold (CM) or regular (non-CM) font. **(b)** the number of developmentally regulated genes in each of the nine species (left) and the number of gene families in which these genes grouped (shared by ≥7 species, right). Groups of gene families that are developmentally regulated in ≥3 Agaricomycotina or ≥4 Pezizomycotina are also shown. **(c)** developmentally regulated gene families grouped by evolutionary conservation and history.

To identify developmentally relevant genes, we used publicly available fruiting body transcriptomes of 5 Pezizomycotina (*33*–*36*) and 4 Agaricomycotina (*37*–*39*) species, with which it was possible to quantify gene expression across 2–13 developmental stages. Based on expression dynamics, we detected 2645–9444 developmentally regulated genes in the nine species (Fig. S1), corresponding to 19.8–66.3% of the proteome. The identified developmentally regulated genes contained 26.9-97.6%, 4.6-69.2% and 92.7% of known developmental genes of *Neurospora, Aspergillus* and *Coprinopsis,* respectively (Note S1, table S2), consistent with previous studies of fruiting body development (*23, 28*–*30, 40*). In a broader dataset of 19 species (see Methods) developmentally regulated genes fell into 21,267 families, of which we focused on that ones that showed conserved developmental regulation in the majority of species (Fig. 1/b). We identified 1,026 gene families that were developmentally regulated in ≥75% of the species in either or both clades, resulting in 314, 273, and 439 families that have a conserved developmental expression in ≥7 of 9 Dikarya, ≥4 of 5 Pezizomycotina and ≥3 of 4 Agaricomycotina species, respectively (table S3). We hereafter focus on these families because these are most likely to have been developmentally regulated also in the most recent common ancestor of Agaricomycotina and that of the Pezizomycotina.

### Widespread parallel co-option of developmental families

We analyzed the origin of the genetic bases of CM by reconstructing the evolution of developmental gene families along the phylogeny. Of the 1,026 conserved developmental families, 560 predate the origin of CM, consistent with their co-option for multicellularity-related functions, while 297 and 169 families are taxonomically restricted to the Agarico-and Pezizomycotina, respectively (Fig. 1/c). Of the 560 ancient families, 314 (56.1%) are developmentally regulated in both the Agarico-and Pezizomycotina, indicating parallel co-option for fruiting body development. The remaining 246 families can be divided into those that have homologs in only one CM clade and were lost in the other (24.5%, 137 families) and those that have homologs in both CM clades but are developmentally regulated only in one (19.5%, 109 families), consistent with clade-specific co-option. The frequency of clade-specific co-option is low, with 42 and 67 families in the Agarico-and Pezizomycotina (7.5% and 12.0%), respectively. The observation of limited clade-specific, but widespread parallel co-option suggests that gene families with suitable properties for CM rarely escaped integration into the genetic toolkit of CM. It also agrees with genes suitable for a given phenotype being rare and thus mostly being recruited under similar selection regimes (*41*).

The observed distribution of developmentally regulated gene families is consistent with two hypotheses. Families with clade-specific phylogenetic distribution or clade-specific developmental expression conform to expectations under the simplest model of convergent evolution at the phenotypic level: two independent gains of CM in the Agarico-and Pezizomycotina. Similarly, shared developmentally expressed families could have been independently co-opted in the two CM clades. However, this set of families could also encode plesiomorphic functions that were present in the Dikarya ancestor and were independently integrated into CM in the Agarico-and Pezizomycotina. Some of those might have served as precursors to CM (e.g. as traits linked with asexual development (*42*)), which could have predisposed lineages for evolving CM independently, leading to a higher likelihood for phenotypic convergence (*23, 43, 44*).

Although the Dikarya ancestor most likely did not have fruiting bodies (see Fig. 1/a), we reasoned that its ancestral gene complement could reveal whether evolutionary predisposition is a reasonable hypothesis to explain CM in fungi. The Dikarya had 989 genes in the 314 shared developmental families (Fig. 2), which we functionally characterized by examining *Saccharomyces cerevisiae* orthologs. Analyses of Gene Ontology terms revealed an enrichment of genes for the regulation of growth, filamentous growth in response to starvation, transmembrane transport, cell communication, gene expression regulation and carbohydrate metabolism, reminiscent of general functions required for fungal development (Fig. S2). We find that several gene regulatory circuits, including ones involved in sexual reproduction, mating partner recognition, light, nutrient and starvation sensing, fungal cell-wall synthesis and modification, cell-to-cell signaling and morphogenesis have been present in the Dikarya (and even earlier), suggesting that these might have provided a foundation for the evolution of fruiting bodies.

**Figure 2.**
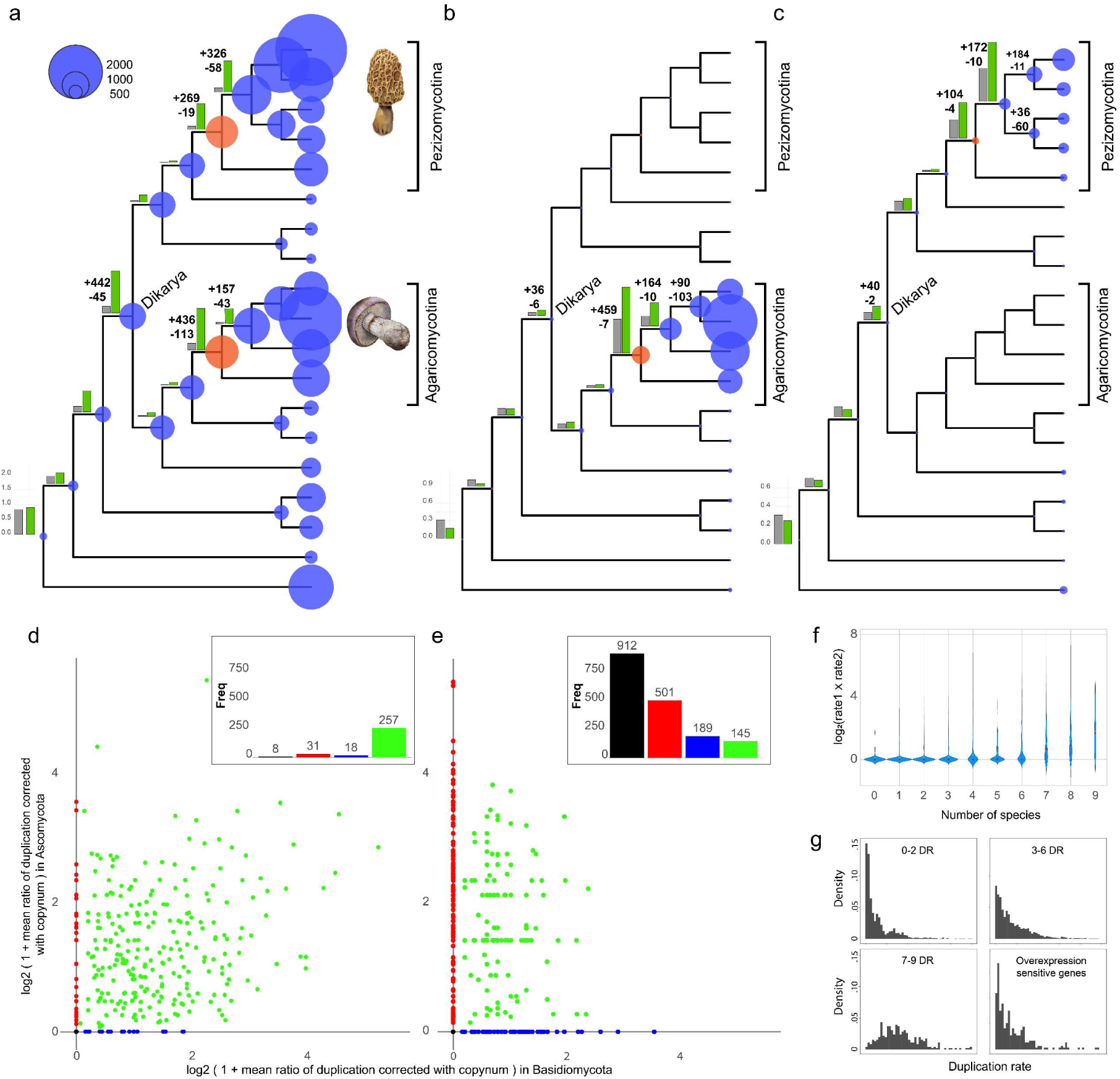
Convergent expansion of developmentally regulated gene families in independent complex multicellular fungi. **(a-c)** Reconstructed copy number evolution of 314 shared developmentally regulated gene families (a), 439 families with Agaricomycotina-specific developmental expression and (c) 273 families with Pezizomycotina-specific developmental expression. Bubble size proportional to the number of reconstructed ancestral gene copies across the analyzed families. Numbers next to internal nodes denote the number of inferred duplications and losses. Bar graphs show genome-wide duplication rates (grey) versus duplication rates of the depicted developmental families (green). Inferred gains of CM are indicated by red bubbles. **(d-e)** scatterplot of Agarico-and Pezizomycotina duplication rates across 314 shared developmentally regulated gene families (d) and 1747 families containing ≤2 developmentally regulated species (e). Black, red, blue and green denote families with no duplications, Ascomycota specific-, Basidiomycota specific-and parallel duplications, respectively. Bar diagrams show the number of gene families in each category. **(f)** correlation between the extent of convergence and the number of species contributing developmentally regulated genes to a family. **(g)** the distribution of gene duplication rates across gene families containing developmentally regulated genes from ≤2, 3-6 and ≥7 species and families in which dosage effects constrain duplications rates (*45*).

### Gene family expansions correlate with the origins of CM

To obtain a higher-resolution picture of the evolution of developmental gene families, we reconstructed gene duplications and losses in the Agarico-and Pezizomycotina and in other parts of the fungal tree. We found characteristic expansions of developmentally regulated gene families in CM Agarico-and Pezizomycotina, but no or significantly smaller expansions in other families (Fig. 2/a, fig. S3). Across the 314 shared families, we inferred a net expansion (duplication minus loss) of 323 and 250 genes in the MRCA of the CM Agarico-and that of CM Pezizomycotina, respectively, indicating that the origin of these groups coincided with significant expansions in developmentally regulated gene families. In the MRCA of Dikarya we found 442 duplications and 45 losses. The observed gene family expansions were driven by increased gene duplication rates, with loss rates remaining approximately constant (Fig. S4). A 6.3 to 8.1-fold higher rate of expansion was found in the 314 shared developmental families compared to other families shared by ≥7 of the nine species, indicating that CM-related gene families are one of the most expanding group in the fungal genomes examined here.

Gene families with a developmental expression specific to the Agarico-(439 families) or Pezizomycotina (273 families) show higher duplication rates (1.93–1.95-fold) and substantially expanded in their respective clades, but not in the other CM subphylum or in non-CM species (Fig. 2/bc). Of the Agarico-and Pezizomycotina the latter showed a more gradual expansion of gene families: we reconstructed 104 and 162 net gains in the MRCA of Pezizomycotina and that of Sordariomycetes, respectively. Interestingly, we inferred relatively few (73) duplications along the branch leading to *Pyronema*, a representative of apothecium-forming Pezizomycotina. This species has 287 genes in the 273 Pezizomycotina-specific families, whereas other species have 401–849 genes. Given that *Pyronema’s* fruiting bodies probably reflect the ancestral morphology (apothecium) in the Pezizomycotina (*46*–*49*), these figures could indicate that the developmental gene repertoire of *Pyronema* resembles the ancestral condition in the Pezizomycotina.

### Convergent expansions in shared developmental families

We next asked whether the expansions observed in two CM clades were composed of expansions of the same gene families (i.e. convergent) or composed of expansions of differing gene families. We calculated subphylum-specific gene duplication rates in the Agarico-and Pezizomycotina, which we plotted against each other as shown on Fig. 2/d. Of the 314 shared developmental families, 257 (81.8%) showed parallel expansions in the Agarico-and Pezizomycotina. In contrast, only eight (2.5%) showed no duplications in either class and 49 (15.6%) showed duplications in only one. If families that likewise contained at least seven species but developmentally regulated genes from up to 2 species (1747 families) were considered, the pattern was the opposite: only 145 (8.3%) showed parallel duplications in the Agarico-and Pezizomycotina, 1602 (91.7%) showed no duplications or in only one of the subphyla (Fig. 2/e). Convergent expansion in the 314 shared developmental families correlates with the number of species represented by developmentally regulated genes in the family (Fig. 2/f). Convergence in shared developmental families is significantly more abundant than in any other combination of gene families we tested (*P* < 1.2×10^−9^, Fisher’s exact test), including controls for gene family size and the number of developmentally regulated proteins per family, among others (table S4, fig. S5). The accelerated duplication rate in these families differs considerably from that of other families (table S4), or families with constraint on duplication imposed by a fitness cost of increased dosage (*45*) (Note S3, Fig 2/g), collectively suggesting that the convergent gene family expansions were driven by positive selection. Families with clade-specific developmental expression, on the other hand, did not show signs of convergent expansion (Fig. S6). We note that further convergent expanding can certainly be found in groups with <7 species, which renders our estimate of convergence conservative.

We also examined the extent of convergence in amino acid sites among CM Agarico-and Pezizomycotina, using approaches that incorporate null models (*50*) proposed in response to previous criticisms of published cases (*51*–*53*). We found 129 families in which convergent shifts in amino acid preference are significantly enriched relative to control analyses (Fig. S7, note S2). Developmentally regulated genes are enriched in 28 of these (Note S2), including genes related to cell division and DNA repair, splicing, ergosterol biosynthesis, among others (see table S5). Nevertheless, the extent of convergence in CM clades was overall similar to that observed in other combinations of clades (note S2), which could indicate that the extent of amino acid convergence in CM is either not outstanding in CM fungi or that other, unknown traits drove convergence also in non-CM clades.

The observed extent of convergence in gene family co-option and expansion exceeds expectations based on previous predictions (*54*) or examples (*55*–*60*) at this phylogenetic scale. In closely related populations of the same species or sister species, evolution works with the same standing genetic variation, providing for a higher incidence of (potentially non-adaptive) genetic parallelism (*3, 61*). Because the probability of repeated recruitment of genes declines with phylogenetic distance, much less convergence is expected among distantly related clades that diverged in the architecture of gene regulatory networks, even if the genes themselves are conserved. Molecular clock estimates suggest that the Agarico-and Pezizomycotina diverged >650 million years ago and their ancestors existed >270 myr after the Dikarya ancestor (*62, 63*). Because of this deep divergence, the extent of parallel co-option and convergent diversification of developmental families in CM Agarico-and Pezizomycotina is not explainable by phylogenetic proximity or neutral processes alone.

## Discussion

Our genome-wide analyses revealed extensive parallel co-option of ancient genes and convergent gene family expansions in two complex multicellular clades of fungi. We observed molecular convergence in hundreds of gene families, with ∼82% of shared developmentally regulated families showing convergent expansions. Several recent studies suggested that molecular convergence may be widespread in nature (*51, 58, 60*). However, while most previous examples were restricted to a few genes (*55, 60, 64, 65*) or to closely related species (*61, 66*), our results suggest that molecular convergence can be pervasive in clades separated by >650 million years of evolution and can affect hundreds of gene families.

The repeated emergence of CM in fungi suggests that evolution can be deterministic, which, in the context of Gould’s famous thought experiment (*67*), means that if we replayed life’s tape, CM would again evolve in fungal clades. This predictability has been attributed to shared genetic variation (*10*), similar selective regimes (*5, 8, 11*) or constraints on the array of acceptable changes (*8*) and on how novelty arises (*4, 8*). A special case of bias in the emergence of novelty is when genes with suitable biochemical properties are available in ancestral species and can easily be co-opted for the same functionalities. Our results are compatible with the scenario that ancient fungi have been predisposed for evolving CM by a rich repertoire of genes in the Dikarya ancestor that are used by extant species for CM-related functions. Predisposed lineages are more likely to show phenotypic convergence (*8, 23, 43, 68*), purely because of the availability of genetic tools that can be recruited for the same functions. It follows that if predisposition indeed happened, then the repeated evolution of CM is not as surprising as it may seem, given the availability of genetic mechanisms that are crucial for the evolution of such multigenic phenotypes.

Already Haldane speculated that similar phenotypes emerge not only as a result of similar selection pressures but also as a result of shared genetic biases (*69*). There is probably a finite number of ways by which CM-associated functions, such as cell adhesion, communication or differentiation can evolve, which explains why the same gene families were co-opted and started diversifying in complex multicellular Agarico-and Pezizomycotina. Our study provides an example on how the genomic repertoire may channel phenotypic evolution towards similar solutions and how this can lead to extensive genetic convergence even at large phylogenetic scales. Such genetic biases on phenotypic evolution suggest that the tireless tinkering of evolution is not only limited by the environment, but also by the genetic ingredients at hand.

## Acknowledgements

We acknowledge inspiring discussions of this topic in the Fungal Genomics and Evolution Laboratory (Szeged, Hungary).

## Author contributions

ZM and LGN conceived the study. ZM, WZ, ANP, JPT, BH and BB analyzed data. ANP analyzed transcriptomic data, WZ, JPT and ZM evaluated developmentally regulated genes, ZM, BH and BB reconstructed gene family evolution. ZM, KK and BP evaluated adaptivity of gene family expansions. ZM, BP and LGN wrote the paper. All authors have read and commented on the manuscript.

## Data and materials availability

All data is available in the main text or the supplementary materials.

## Supplementary Materials

### Materials and Methods

#### Bioinformatic analysis of RNA-Seq data

We downloaded publicly available transcriptome data related to different developmental stages of nine fruiting body (FB) forming species from the Pezizomycotina (*Fusarium graminearum, F. verticillioides, Sordaria macrospora, Neurospora crassa, Pyronema confluens*) and the Agaricomycotina (*Armillaria ostoyae, Coprinopsis cinerea, Hypsizygus marmoreus, Flammulina velutipes*, table S1). We ran a quality check on raw fastq files using fastqc v0.11.5 (*70*) and trimmed the adapter sequences and low-quality bases with trimmomatic v0.36 (*71*). Next, we used kallisto v0.43.1 (*72*) to quantify the abundance of transcripts for each stage. Specifically, we utilized the estimated counts from abundance data to calculate Fragments Per Kilobase Million (FPKM) and used it as the quantification metric. As a pre-filter, an FPKM value less than two was considered insignificant. The homogeneity of biological replicates was checked by constructing MDS plots based on overall expression levels.

#### Identification of Developmentally Regulated Genes

The RNA-Seq data comprised 2-13 stages, which was used to identify developmentally regulated genes: those that show at least four-fold change in expression between any two fruiting body stages or tissue types and that show an expression level FPKM > 4. Fold change values were calculated for all biologically relevant pairwise comparisons (for see details fig. S1).

#### Comparative genomic approaches

In addition to the nine above mentioned fruiting body forming species, 10 additional species were included in the analysis for comparative purposes (table S1). This 19 genome dataset was clustered into gene families using OrthoFinder v1.1.8 (*73*) with the default inflation parameter of 1.5 to facilitate interspecies comparison.

For functional annotation of genes and gene families InterProscan search was performed with InterProscan version 5.28-67.0 (*74*) across the 19 fungal proteomes (table S3).

Gene Ontology (GO) enrichment analysis for yeast orthologs was performed using GOrilla ((*75*) http://cbl-gorilla.cs.technion.ac.il/) with *Saccharomyces cerevisiae* as the reference organism, a 10^-3^ P-value threshold and false discovery rate correction for multiple testing. Terms in all three ontologies (Biological process, Cellular component, Molecular function) were considered. Experimentally verified gene function from *Aspergillus nidulans* and *Neurospora crassa* were also considered during the functional annotation of developmentally regulated gene families. We also used known developmentally regulated gene set of *Coprinopsis* from literature to verify the efficiency of our designation.

#### Phylogenetic analyses and ancestral state reconstructions

Altogether 86 clusters were single copy and shared by all 19 species; these clusters were used to reconstruct a species tree. After multiple sequence alignment using the L-INS-I algorithm of MAFFT (*76*) and trimming with trimAL (–gt 0.6) (*77*) sequences were concatenated into a supermatrix and used for phylogenetic reconstruction in raxmlHPC-PTHREADS-SSE3 (*78*). The supermatrix was partitioned by gene and the PROTGAMMAWAG model was used with 100 rapid bootstrap replicates, to estimate branch support.

To reconstruct the evolution of fruiting body formation, maximum likelihood ancestral state reconstructions were performed with the ace (ancestral character estimations) function of the ape R package ((*79*); R Development Core Team, 2018). As the more parametrized ARD (all-rates-different) model didn’t yield significantly greater likelihoods, we used the ER (equal rates) model.

#### Evolutionary history of gene families

To reconstruct the duplication and loss events of gene families across the species tree, the COMPARE pipeline (*43*) was used. For this analysis gene trees were reconstructed for clusters containing at least four proteins (9686 clusters). Sequences in each cluster were aligned using the L-INS-I method of MAFFT and trimmed with trim-AL (-gt 0.2). Gene tree reconstructions were performed in RAxML under the PROTGAMMAWAG model with 100 rapid bootstrap replicates. Gene trees were rerooted and reconciled with the species tree using Notung 2.9 (*80*) with 80% bootstrap support as the edge-weight threshold for topological rearrangements. After ortholog coding, duplications and losses for each orthogroup were mapped onto the species tree using Dollo parsimony (*43*). The visualization of reconstructed duplication/loss histories and further statistical analyses (Fisher Exact test) were performed with custom R scripts (available from the authors upon request).

To quantify convergent gene family expansions, we filtered families with the following criteria: a) a gene family has genes conserved in ≥7 of the 9 Dikarya species, ≥4 of the 5 Pezizomycotina species or ≥3 of the 4 Agaricomycotina species or b) a cluster has developmental expression conserved in ≥7 of the 9 Dikarya species, ≥4 of the 5 Pezizomycotina species or ≥3 of the 4 Agaricomycotina species.

#### Convergence in gene family expansions

To quantify the level of convergent gene family expansions, subphylum-specific gene duplication rates were compared between the Agarico-and Pezizomycotina. Rates were calculated by normalizing the raw number of inferred duplications for a given node by both the length of the preceding branch and by gene family size, because along longer branches and in larger gene families the probability of duplications is naturally higher. After this correction step, duplication rates were averaged across nodes (gene duplication rate for Agarico-and Pezizomycotina) and plotted using custom R scripts. The numbers of four possible events were recorded: duplications in only one (Agarico-or Pezizomycotina), both or none of the CM clades.

To assess if developmental gene families show more or less convergence than expected by chance, different control groups of gene families were generated and compared using Fisher’s exact test. Control families were always compared to the conserved developmentally regulated cluster set (developmentally expression conserved in ≥7 of the 9 Dikarya species). The first control groups comprised families that similarly contained ≥7 species but only 0-2 (1747 cluster) or 3-6 species (2052 cluster) with developmental expression. Next, to test if gene family size (i.e. number of proteins) impacts convergence, we also generated control groups with similar gene family size distribution but containing less developmentally regulated genes than the 314 shared developmental gene families. A custom R script was used to find a non-developmentally regulated gene family for each of the developmentally regulated gene family with a matching size one by one. If it was not possible to find a gene family with similar size (permitted maximum difference of 10%), the gene family was excluded from the comparison. If there were more than one gene families with the same size, the one with most similar species composition and least developmentally regulated genes (according to number of species represent developmentally regulated genes) was chosen.

#### Detecting convergent amino acid changes

In order to gain insights into amino acid convergence between the Agaricomycotina and Pezizomycotina, we followed Rey et al.’s approach (*50*) to identify convergent shifts in amino acid preference at a given site. Convergence is defined not only as changes to identical amino acids from different ancestral states, but also as changes to amino acids with similar biochemical properties (referred to as convergent shifts in amino acid composition). We identified such shifts across all gene families in the 19 species’ genomes using the model “Profile Change with One Change” (PCOC) (*50*) (downloaded from https://github.com/CarineRey/pcoc on 2018.11.05). We used reconciled gene trees with branch lengths re-estimated with RAxML (raxmlHPC-PTHREADS-SSE3) as input. Each of the most inclusive clades that contained only CM species were designated as phenotypically convergent clade. Automated designation of convergent clades was done using a custom R script, followed by execution of PCOC with default settings. For considering a site as convergent we chose the PCOC model with a posterior probability threshold of 0.8. We performed the analysis on 3799 gene families that contained at least 10 proteins and at least one protein from both the Agaricomycotina and the Pezizomycotina. Three sets of control analyses were run assess the amount of convergence caused by chance events. In control 1, the basal lineages of Ascomycota and Basidiomycota were designated as convergent clades (Ustilagomycotina, Pucciniomycotina, Saccharomycotina, Taphrinomycotina). In control 2, CM Agaricomycotina species were paired with the basal clades of Ascomycota (Saccharomycotina, Taphrinomycotina) while in control 3 CM Pezizomycotina were paired with the basal clades of Basidiomycota (Ustilagomycotina, Pucciniomycotina). This resulted in three control analyses in which CM is not shared by clades designated as convergent (note however, that other traits might be). The numbers of detected amino acid sites showing convergent shifts in each gene family were recorded and correlations between CM and control groups were evaluated with a Pearson correlation test. We also compared these values after correction by branch lengths between or in the designated clades to avoid the effect of divergence (i.e. branch length) on the amount of amino acid changes.

We assumed that gene families which contain more convergent amino acid sites in CM lineages than in non-CM clades might be involved in the shaping of convergent phenotypes. For identification of these gene families, a linear model was fit to predict the number of convergent sites between CM clades from the corresponding values of non-CM clades (control 1). Gene families with more convergent sites than the upper limit of 95% prediction interval of the linear model were considered as displaying significant number of convergent sites in CM clades.

### Supplementary Text

#### Comparison of developmentally regulated genes with known developmental genes

To validate our approach to identifying developmental genes, we compared our sets of developmentally regulated genes to suites of genes known to be involved in fruiting body development in *Aspergillus nidulans, Neurospora crassa* and *Coprinopsis cinerea*. First we compared our dataset to that of (*28*). They ranked genes with the largest evolved difference in gene expression change across perithecium development of 5 Ascomycota species (from *Fusarium* and *Neurospora*). Of the 41 genes they found to have an aberrant phenotype in sexual development of either *Neurospora crassa* or *Fusarium graminearum*, our analysis we identified 40 (97.6%, table S2). Another study (*81*) identified 234 genes causing phenotypic changes during fruiting body formation when knocked out in Neurospora, of which 63 were detected as developmentally regulated (26.9%) in our study (table S2). Secondly, we examined homologs of genes involved in the initiation and coordination of sexual reproduction of *Aspergillus nidulans* (84). Homologs of these genes were identified in our nine species based on best BLAST hits (e-value < 10^-6^). Depending on the species, 4.6-69.2% (table S2) of homolog genes were developmentally regulated in our dataset. The lowest ratio is found in *Pyronema confluens* where only two mixed stage (both samples contains vegetative mycelium) were sampled for transcriptomic analysis (*35*). We also collected genes from the literature with a proven role in fruiting body development of *Coprinopsis cinerea*. Our identified developmentally regulated genes contained 92.7% of known developmental genes of *Coprinopsis* (table S2).

#### Convergent amino acid shifts

Altogether 3541 gene families were analyzed with PCOC (*50*) to identify convergent amino acid shifts related to the convergent phenotype of complex multicellularity (CM). When CM species were designated as convergent clades the gene trees of families had on average 2.52±1.65 (max: 41) convergent clades (median: 2) and PCOC identified on average 10.23±10.6 sites as convergent (median: 7, max: 117). In the case of the control comparison consisting of basal, non-CM lineages of the Asco-and Basidiomycota, the gene trees had on average 4.03±2.08 (max: 66) convergent clades (median: 4) and on average 10.66±11.45 sites were detected as convergent (median: 8, max: 123). The numbers of amino acid sites identified by the PCOC model were compared between CM clades and control groups in each of the gene families in order to decide whether the number of sites in CM species is higher than that in the control clades, indicating adaptive amino acid convergence related to CM. The number of detected amino acid sites showed strong correlation (Pearson r > 0.92, p-value < 2.2 ×10^-16^) between the CM and control comparisons (Fig. S8), indicating similar levels of amino acid convergence in CM and non-CM clades. This could be explained two ways. First, it is possible that there isn’t more amino-acid convergence in relation with CM than background levels. Alternatively, unmeasured traits in the control clades could be associated with similar amounts of adaptive amino acid convergence as CM, explaining their similar levels in the two sets of clades. Despite different normalizations (with branch length between or in the designated clades) (Fig. S8), the results did not generally support a higher-than-expected contribution of amino acid convergence to the independent emergence of CM. However, 129 gene families (3.64% of all examined) had more convergent sites than the upper border of prediction intervals of a linear model, in comparison with the control analysis (R^2^= 0.85, p-value < 2.2 ×10^-16^). From these 129 gene families only 28 contain more developmentally regulated proteins than expected by chance (table S5). One of them (OG0002402) is homologous with the *Aspergillus nimO/AN1779*, protein kinase, contained 7 developmentally regulated genes (out of 10 genes in total), and 25 convergently evolved amino acid site between the two subphyla (in contrast: 13-22 in control groups, fig. S9). This gene family might be an example of amino acid convergence related to fruiting body formation.

#### Gene families with constraint

We used ranked gene lists of *S. cerevisiae* from gene overexpression assays of Sopko et al. (*45*), who analyzed and scored (toxicity score from lethal to wild type: 1-5) the growth rate of strongly overexpressing each of 5917 genes in *Saccharomyces*. These overexpression sensitive genes were identified in our dataset using BLAST with the *Saccharomyces* strain S288C as a query (downloaded from https://downloads.yeastgenome.org/sequence/S288C_reference/orf_protein/). After the detection of best hits for each query protein, and an 80% identity filter, we could identify unequivocally 3907 gene families in which all members of the family had the same toxicity score. In 718 cases members of the gene families had different toxicity scores, in such cases we accepted those toxicity scores which were supported by the most BLAST hits, or we took into account the largest score. Finally, the duplication rates of 296 gene families with a toxicity score <3 (most sensitive for gene overexpression in *Saccharomyces*) were compared to the duplication rates of conserved developmental gene families (containing ≥7 of the 9 CM species). In this comparison, the duplication rate of a gene family was averaged across all nodes of the tree and calculated by normalizing the raw number of inferred duplications for a given node by both the length of the preceding branch and gene family size.

---

**Fig. S1.**
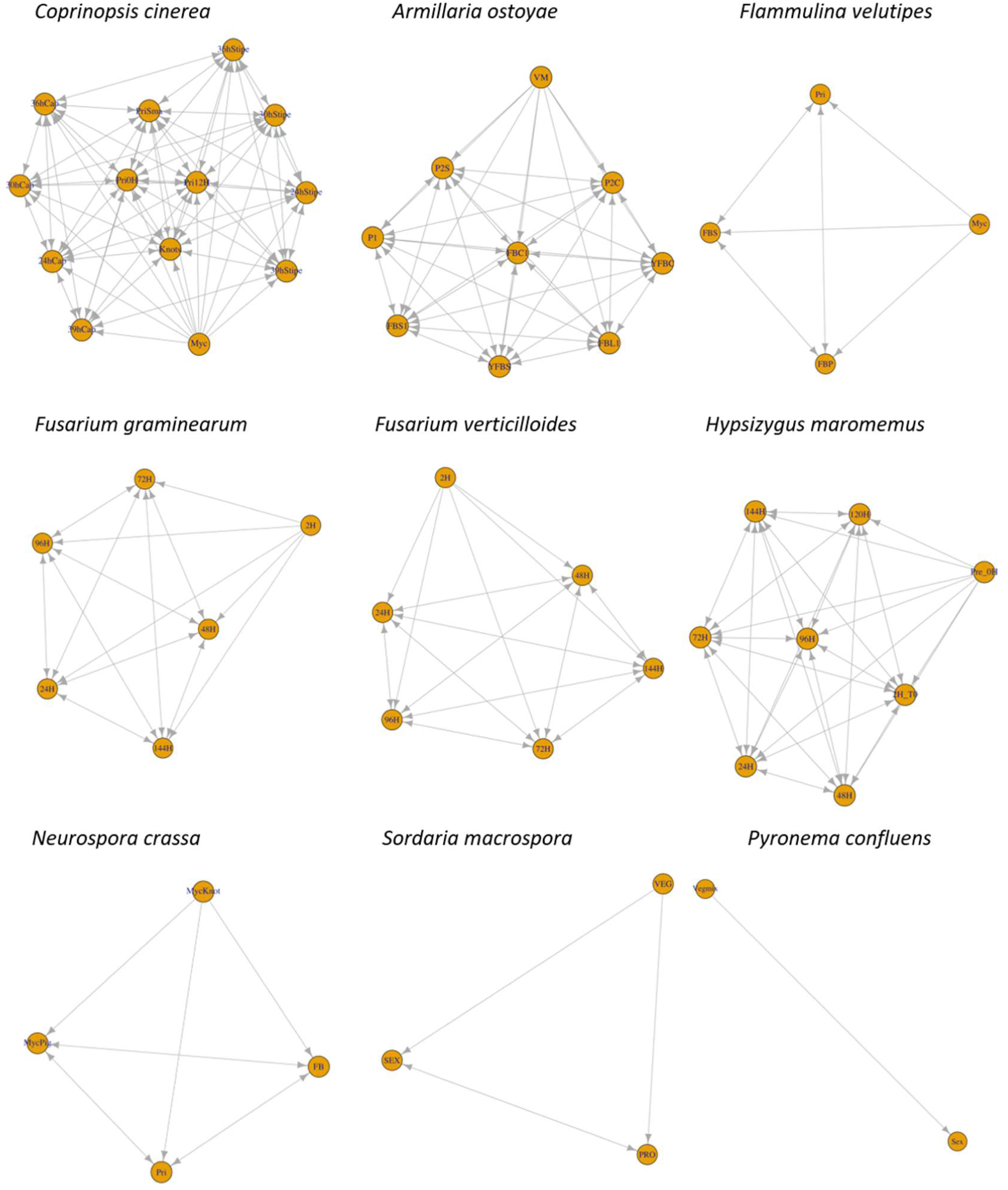
Stage-wise comparisons used for identifying developmentally regulated genes, for each of the nine species. Developmental stages (nodes) and allowed comparisons (arrows) for the identification of developmentally regulated genes are shown. Arrows pointing in only one direction represent unidirectional comparisons, which were done in order to exclude genes showing highest expression in vegetative mycelium and no dynamics later on. Comparisons between tissue types were only allowed within the same developmental stage. H = hour, Pri = P = Primordia, S = Stipe, FB = Fruiting body, Y = Young, VEG= VM = Myc = Mycelium, C = Cap, L = Lamellae.

**Fig. S3.**
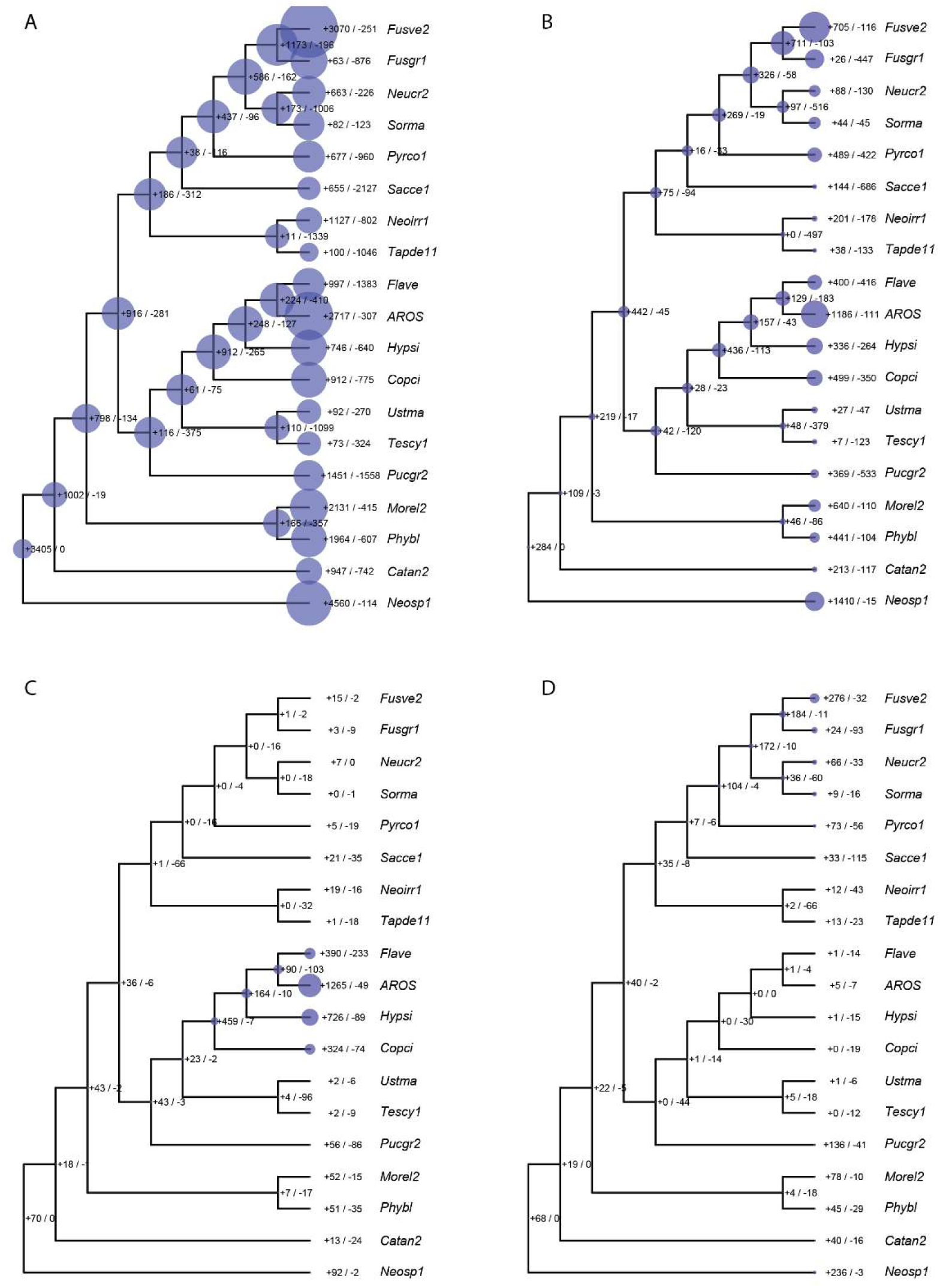
Convergent expansion of developmentally regulated gene families in independent complex multicellular fungi. (A-C) Reconstructed copy number evolution of (A) 4113 shared gene families, (B) 314 shared developmentally regulated gene families, (C) 439 families with Agaricomycotina-specific developmental expression and (D) 273 families with Pezizomycotina-specific developmental expression. Bubble size proportional to the number of reconstructed ancestral gene copies across the analyzed families. Numbers next to internal nodes denote the number of inferred duplications (+) and losses (-).

**Fig. S4.**
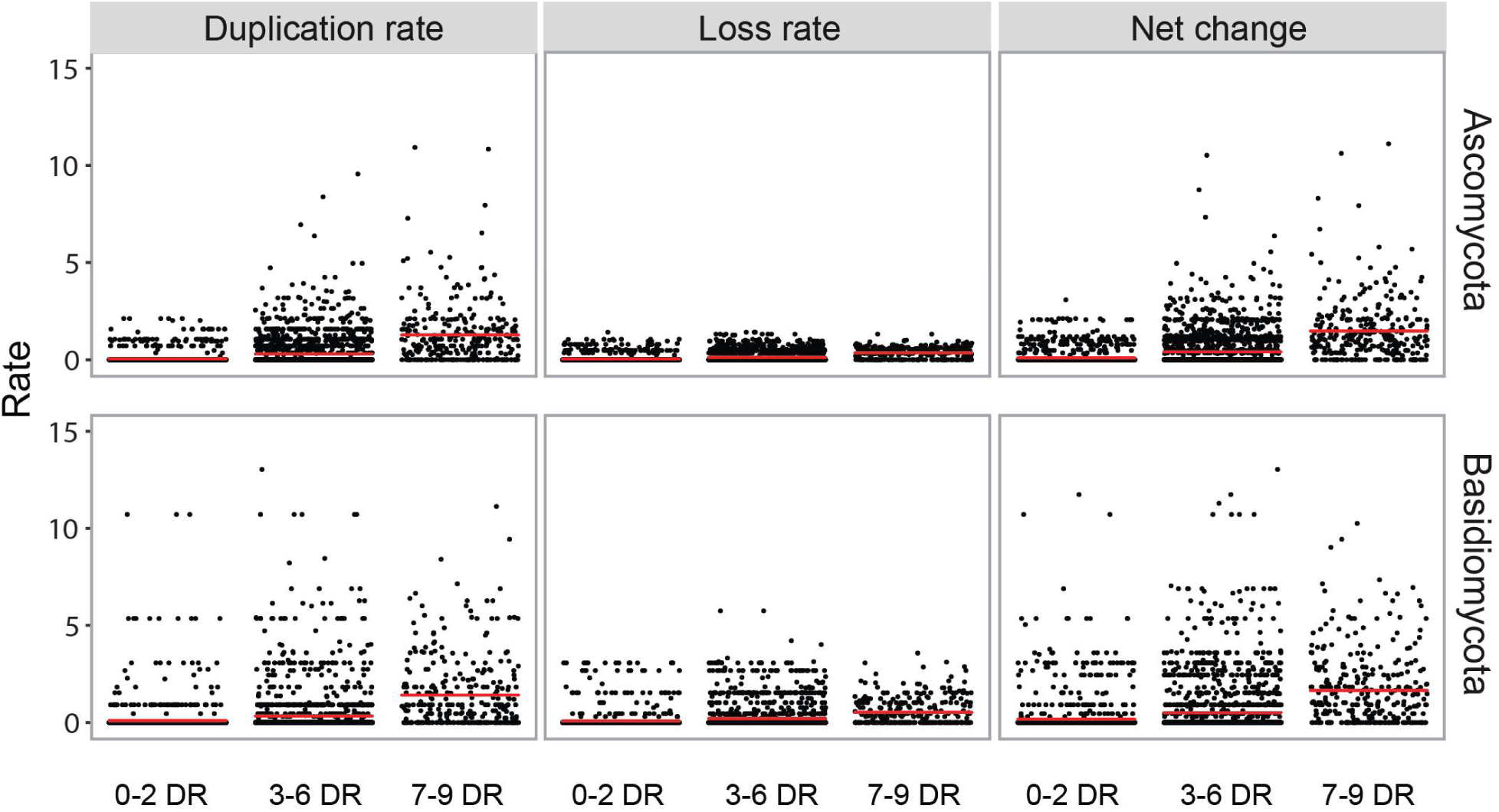
Gene duplication, gene loss and net gene family expansion rates across gene families containing developmentally regulated genes from ≤2 (0-2DR), 3-6 (3-6DR) and ≥7 (7-9DR) species. Red lines mark mean values.

**Fig. S5.**
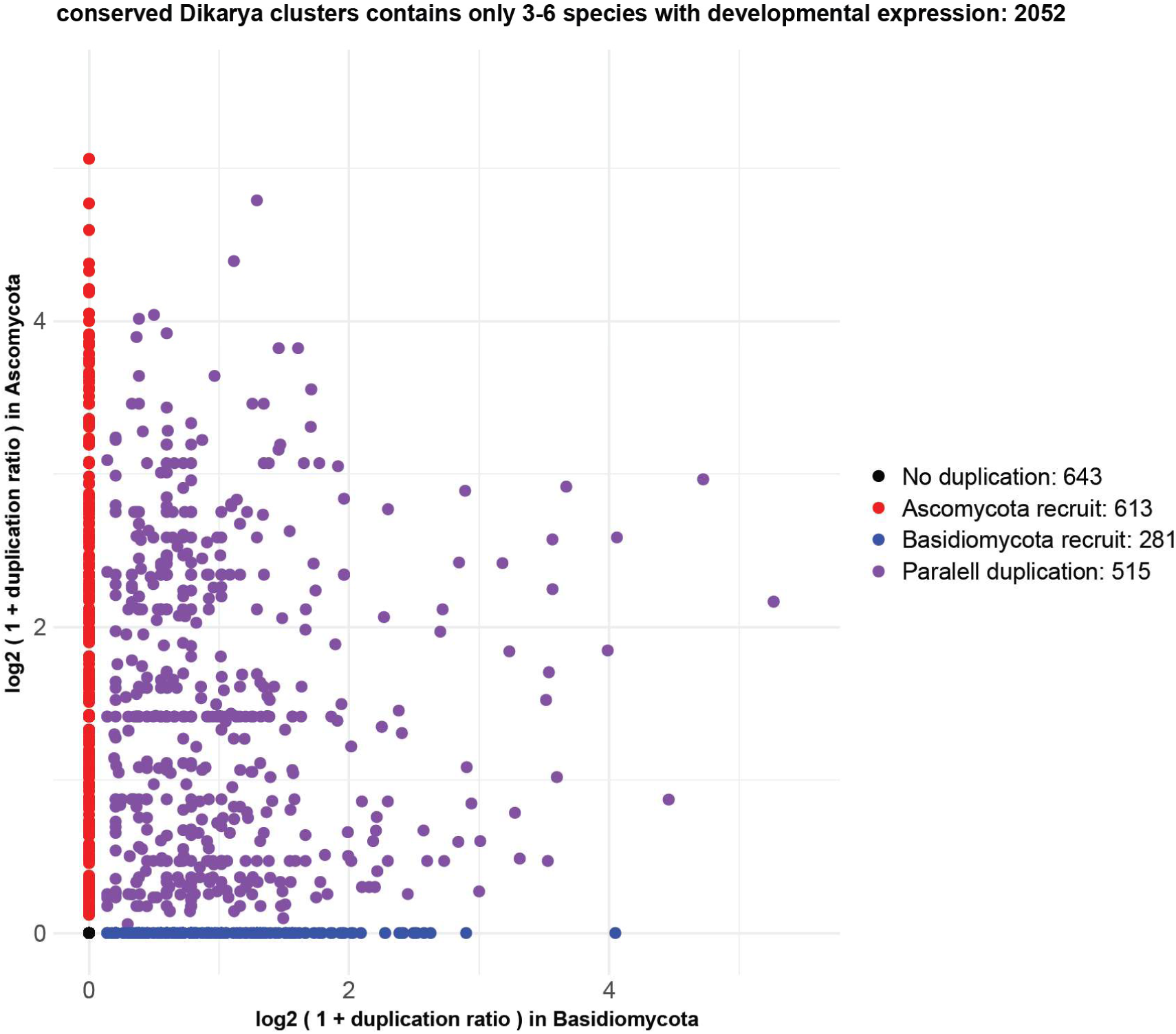
Convergent gene family expansions in conserved Dikarya families containing 3-6 species with developmental expression. Each dot represents a family, while the x and y axes represent the duplication rate in the Agaricomycotina and Pezizomycotina, respectively.

**Fig. S6.**
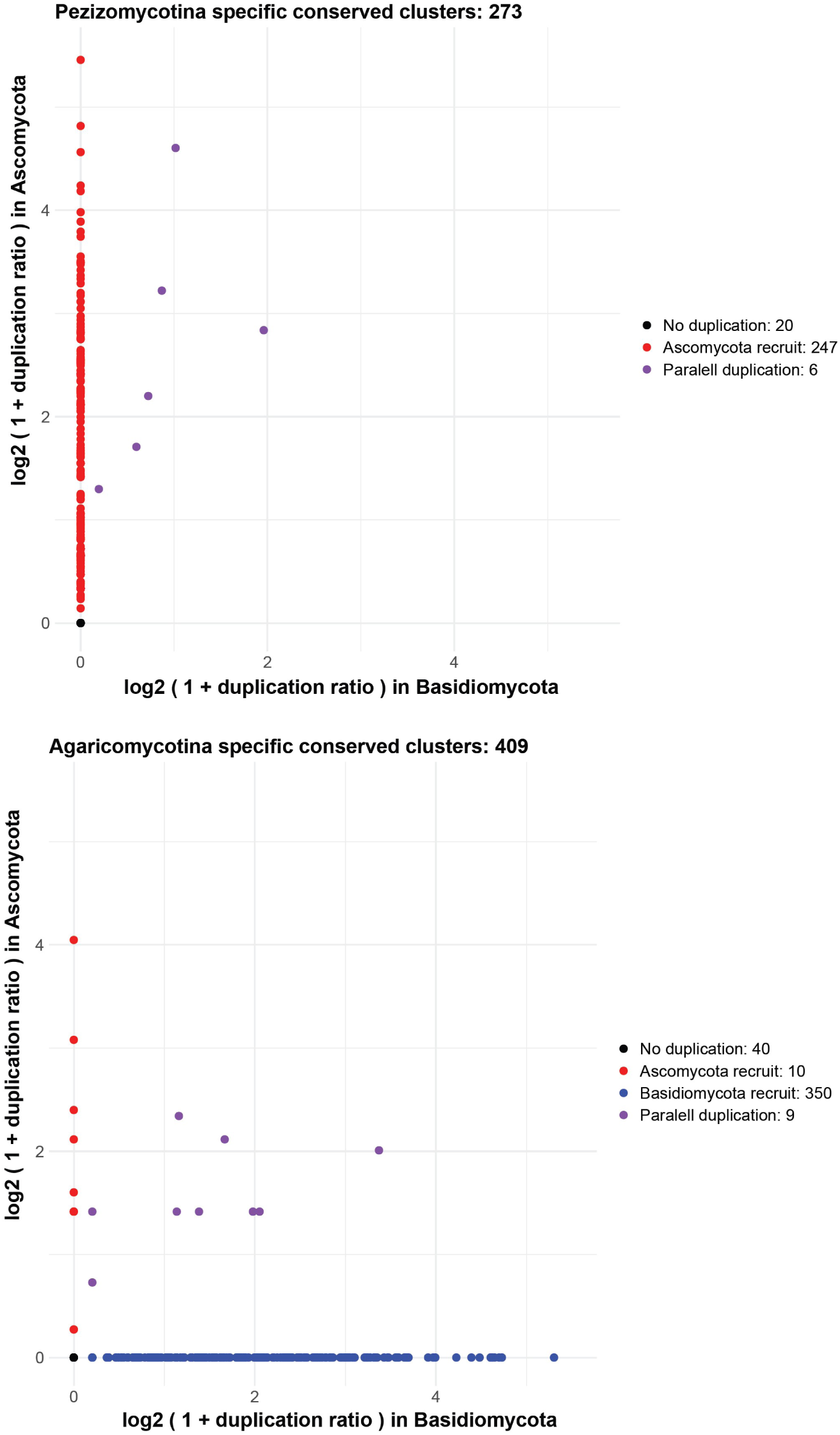
Convergent gene family expansions in gene families with developmental expression specific to the Pezizomycotina (top) or the Agaricomycotina (bottom). Each dot represents a family, while the x and y axes represent duplication rate in Agaricomycotina and Pezizomycotina, respectively.

**Fig. S7.**
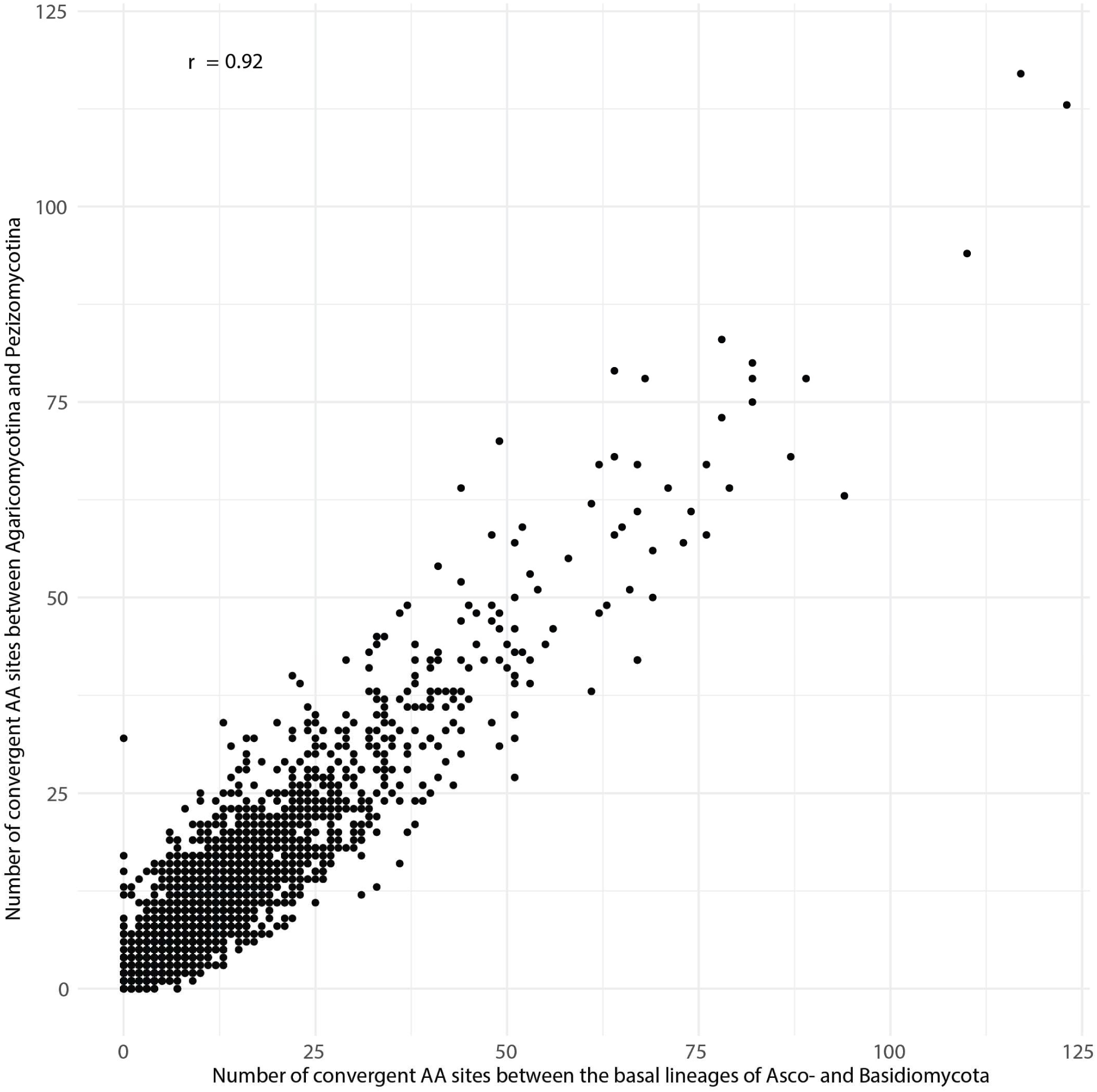
Correlation between the numbers of convergent amino acid (AA) sites identified by the PCOC model in CM clades and in basal Asco-and Basidiomycota clades, as controls groups, across 3541 gene families. Each dot corresponds to a gene family tested for amino acid convergence. The y-axis represents the number of convergent AA sites detected among the Agaricomycotina and Pezizomycotina, while the x-axis represents a control estimate basal lineages of Asco-and Basidiomycota. r denotes Pearson correlation coefficient.

**Fig. S8.**
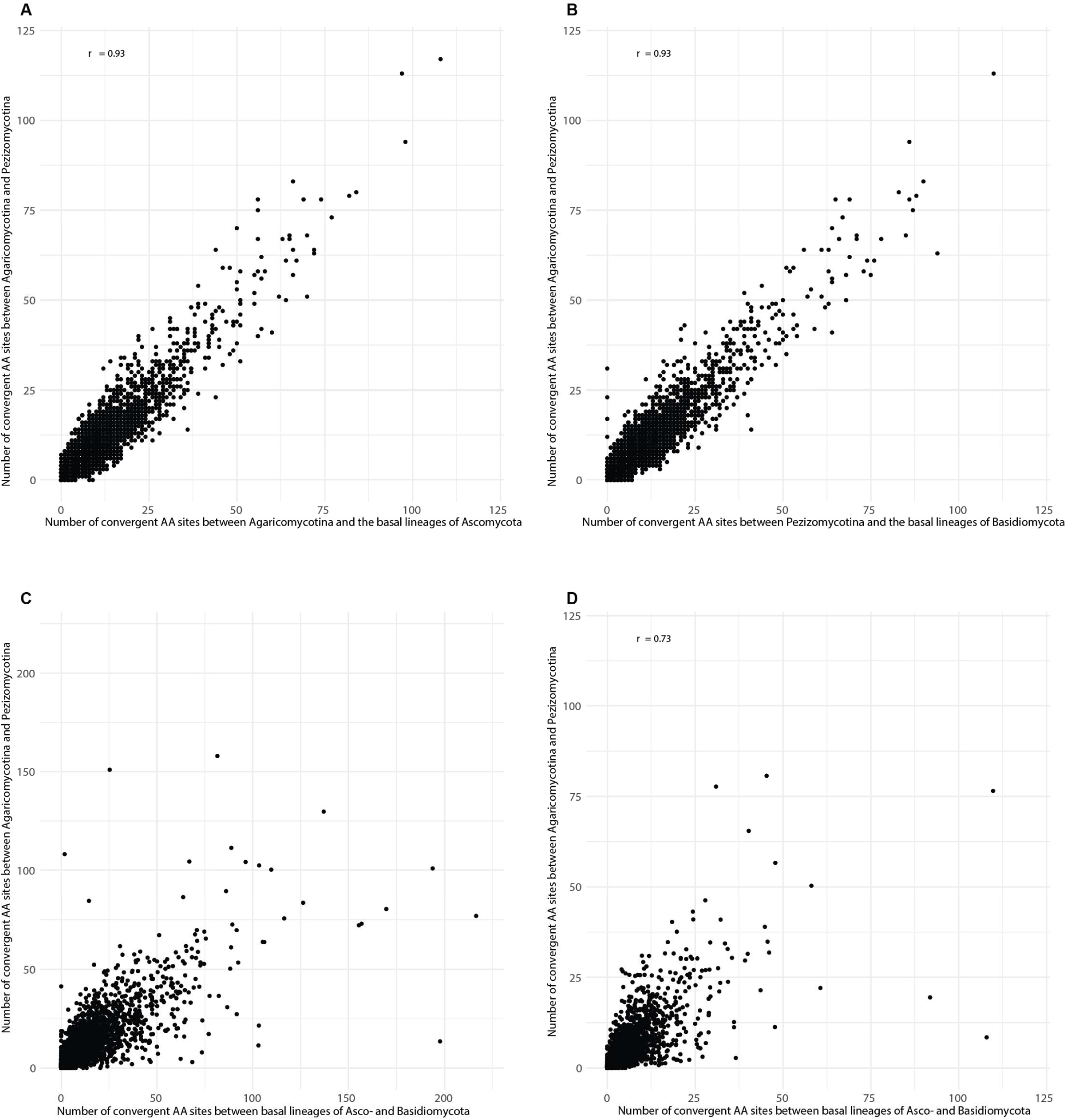
Numbers of convergent amino acid (AA) sites identified by the PCOC model. In each scatterplot the y-axis represents the number of convergent AA sites detected among the fruiting body forming clades (Agaricomycotina and Pezizomycotina), while the x-axis represents a control estimate: A) Agaricomycotina paired with the basal lineages of Ascomycotina (Saccharo-and Taphrinomycotina) B) Pezizomycotina paired with the basal lineages of Basidiomycota (Ustilago-and Pucciniomycotina). C) basal lineages of Asco-and Basidiomycota normalized with the branch length in convergent lineages; D) basal lineages of Asco-and Basidiomycota normalized by the patristic distance between the crown nodes of convergent lineages. r denotes Pearson correlation coefficient.

**Fig. S9.**
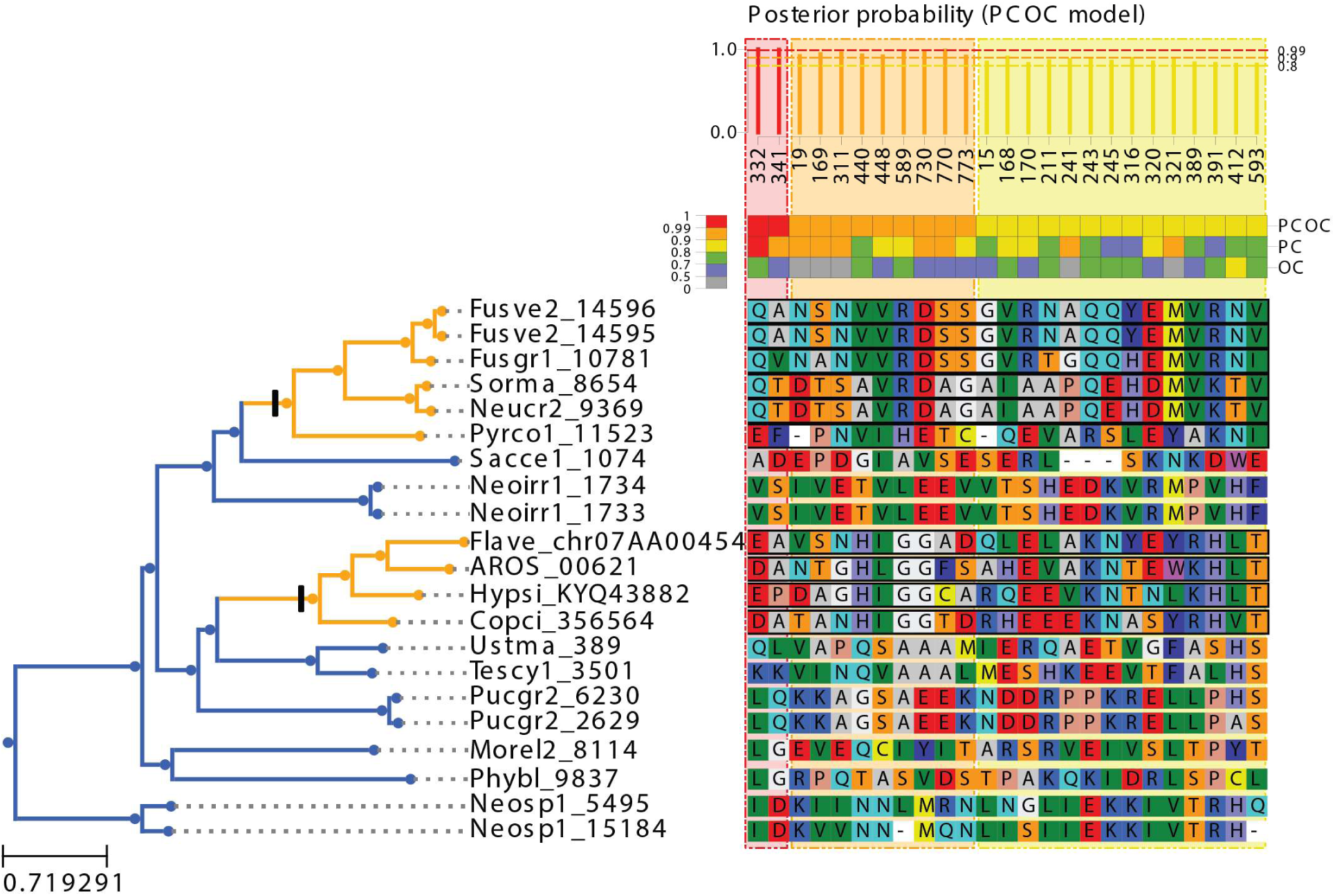
An example of convergent amino acid shifts in the gene family containing *Aspergillus nimO/AN1779* homologs (OG0002402) identified using PCOC. Only sites where the posterior probability of being convergent was above 0.8 are shown. In the gene tree (left side) clades showing convergent phenotypes (fruiting body formation) are highlighted in orange.

**Table S1. (separate file).** List of 19 species used in comparative genomic analyses

**Table S2. (separate file).** Comparisons developmentally regulated genes to homologues with known developmental roles in model organisms of fungal CM.

**Table S3. (separate file).** List of 1026 gene families with conserved developmental expression

**Table S4. (separate file).** Statistical comparisons of the amount of parallel duplication in developmental versus control gene families.

**Table S5. (separate file).** Gene families with significantly more amino acid shifts than expected by chance.

**Fig S2. (separate file).** Gene Ontology analyses for 314 conserved developmental gene families

