## Supplementary material for "Unmatched level of molecular convergence among deeply divergent complex multicellular fungi": Suppl_Fig_2

**Fig. S2.**

Gene Ontology terms of 314 conserved developmental gene family. Despite the much-reduced repertoire of Saccharomyces, we chose this species because of the abundance of experimentally verified functional information on genes.

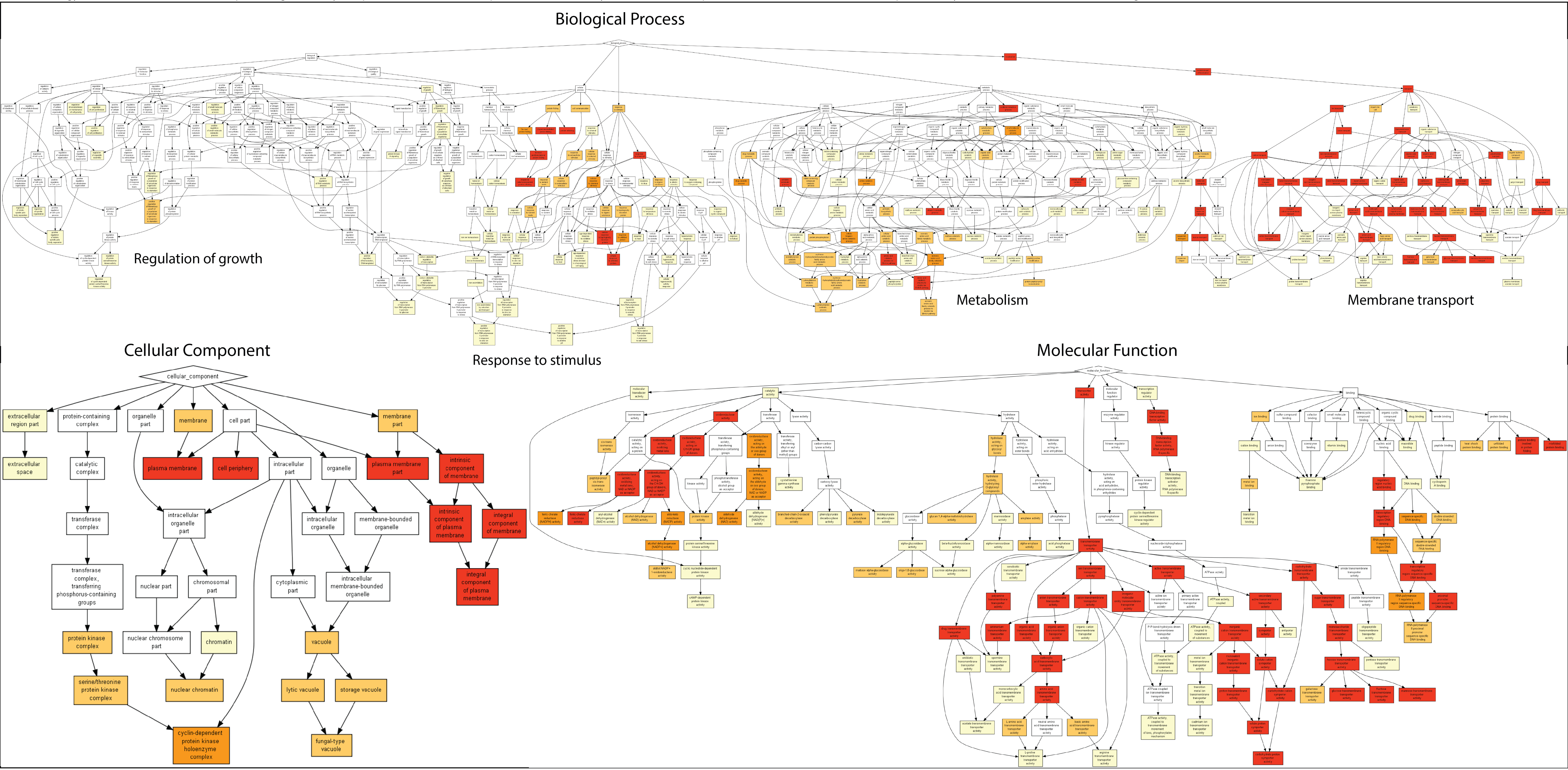
